# *Fusobacterium nucleatum* outer membrane vesicles activate the TNFR1/NF-κB signaling pathway to promote OSCC proliferation

**DOI:** 10.1101/2025.10.31.685796

**Authors:** Zhengrui Li, Ji’an Liu, Rao Fu, Xutao Wen, Xufeng Huang, Divya Gopinath, Ling Zhang

## Abstract

*Fusobacterium nucleatum* (*F. nucleatum,* Fn) has emerged as a significant bacterium linked to cancer. However, the precise mechanisms by which Fn drives the progression of oral squamous cell carcinoma (OSCC) remain obscure. Outer membrane vesicles (OMVs), small vesicular structures secreted by Gram-negative bacteria, are known to contribute to their pathogenicity. This study investigated the role of Fn OMVs in oral cancer progression and the underlying mechanisms.

**Methods:** Metagenomic sequencing was conducted on both oral cancer and control tissues. OMVs were isolated through ultracentrifugation. Cell proliferation assays on two cancer cell lines assessed the impact of Fn and its OMVs. Fn OMVs were administered to mice to evaluate their influence on tumorigenesis. RNA-seq was performed on OMV-treated cells to identify enriched genes, which were validated *via* qPCR and Western blot. Knockdown experiments were executed *in vitro* and *in vivo* to confirm the involvement of Fn OMVs in TNFR1-mediated NF-κB signaling.

**Results:** Fn prevalence was markedly higher in oral cancer tissues compared to non-cancerous samples. *In vitro*, Fn and its OMVs significantly enhanced the growth and proliferation of oral cancer cell lines. Mice treated with Fn OMVs developed larger tumors than those receiving Fn alone. Furthermore, Fn OMVs notably induced TNFR1 expression and activated the NF-κB signaling pathway, as confirmed by RNA-seq, qPCR, and Western blot. TNFR1 knockdown significantly diminished the effects of Fn OMVs on cancer cell proliferation *in vitro* and tumor formation *in vivo*.

**Conclusion:** This study compellingly demonstrates that Fn OMVs are pivotal in OSCC progression by directly activating TNFR1-mediated NF-κB signaling. These findings pave the way for future research into the prevention and treatment of oral cancer.

## Introduction

Oral squamous cell carcinoma (OSCC) ranks among the most prevalent malignant tumors worldwide, posing a significant threat to human health[1, 2]. Despite advancements in treatment, high recurrence rates have led to unsatisfactory long-term survival for OSCC patients[1]. Thus, investigating new molecular mechanisms and therapeutic targets is essential for improving outcomes.

Recent studies have increasingly focused on the link between microbial communities and cancer development, with particular attention on the association between oral microbial dysbiosis and OSCC[3, 4]. Among the common Gram-negative anaerobes in the oral cavity, *Fusobacterium nucleatum* (*F. nucleatum*, *Fn*) has gained considerable interest due to its presence in various tumor microenvironments and its role in tumor proliferation, invasion, and immune evasion[5–8]. *F. nucleatum* interacts with host cells *via* various secretion pathways, modulating signaling and promoting tumor progression[9–11].

Outer membrane vesicles (OMVs) are small structures secreted by Gram-negative bacteria, composed of the bacterial outer membrane and extracellular components[12, 13]. OMVs facilitate communication between bacteria and host cells, carrying biomolecules such as proteins, lipids, and nucleic acids that influence immune responses and cell signaling[12, 13]. However, research on Fn OMVs remains limited, and their potential role in oral cancer is not well understood.

Nuclear factor κB (NF-κB) signaling pathways are crucial in regulating cell survival, proliferation, and inflammation[14]. NF-κB activation has been linked to aggressive cancers, including OSCC[15, 16]. Tumor necrosis factor receptor 1 (TNFR1), a type 1 transmembrane receptor, plays a pivotal role in NF-κB activation and OSCC progression[17–20]. It was hypothesized that Fn OMVs might activate TNFR1, thereby regulating NF-κB signaling and promoting OSCC cell proliferation.

This study aimed to determine whether Fn OMVs promote OSCC proliferation through TNFR1-mediated NF-κB regulation. Results demonstrated that Fn OMVs significantly enhanced OSCC cell proliferation *via* TNFR1 and subsequent NF-κB signaling pathways. This novel finding uncovers a mechanism by which *F. nucleatum* modulates host cell signaling through OMVs, revealing new potential targets for OSCC molecular therapy. Moreover, these results highlight the role of microbial communities in cancer development, offering a new perspective for future treatment strategies. The development of therapies targeting microbial regulatory mechanisms based on these findings may provide new hope for OSCC patients.

## Methods

### OSCC Clinical Sample Collection

The study included patients diagnosed with OSCC from the hospital and healthy volunteers. Patients were selected based on the following inclusion criteria: a confirmed diagnosis of OSCC with no other mucosal lesions in the oral cavity. Exclusion criteria included recent antibiotic use within the past month, prior radiotherapy or chemotherapy, and a history of other malignant tumors. The healthy control group consisted of individuals without periodontal disease or other oral disorders as confirmed during the physical examination. Their exclusion criteria mirrored those of the OSCC group, including recent antibiotic use, previous radiotherapy or chemotherapy, and a history of malignant tumors. From January 2023 to September 2023, 40 OSCC patients and 10 healthy controls were enrolled in the study (Table 1).

**Table 1.** Clinical characteristics of patients with OSCC.

| Group | Normal<br>Control Group | Experimental Group |  |  |  |
| --- | --- | --- | --- | --- | --- |
|  | NC | T1 | T2 | T3 | T4 |
| Mean age | 51.7 ± 12.4 | 62.8 ± 7.8 | 60.2 ± 10.0 | 62.6 ± 8.5 | 65.1 ± 5.0 |
| Male | 5 | 4 | 5 | 5 | 7 |
| Smoking | 5 | 4 | 5 | 6 | 5 |
| Drinking | 5 | 2 | 4 | 3 | 6 |
| Anatomical Site |  |  |  |  |  |
| Tongue | 2 | 5 | 4 | 3 | 2 |
| Buccal | 2 | 2 | 2 | 3 | 2 |
| Floor of mouth | 2 | 1 | 1 | 2 | 2 |
| Gingiva | 2 | 2 | 2 | 1 | 2 |
| Palatine | 2 | 0 | 1 | 1 | 2 |
| Total | 10 | 10 | 10 | 10 | 10 |

Samples were collected using single-use sterile nylon swabs and stored in an oral swab preservation solution to prevent DNA degradation. Intraoperatively, after tumor resection, samples were obtained from within the tumor (experimental group: different stages of tumor progression, Table 2) and from adjacent normal tissue (tumor-negative margin) (control group). All samples were transported to the laboratory within 2 hours of collection and stored at −80°C until analysis. The study received approval from the Ethics Committee of the Ninth People’s Hospital, affiliated with Shanghai Jiao Tong University School of Medicine (SH9H-2023-TK212-1), and all participants provided written informed consent.

**Table 2.**
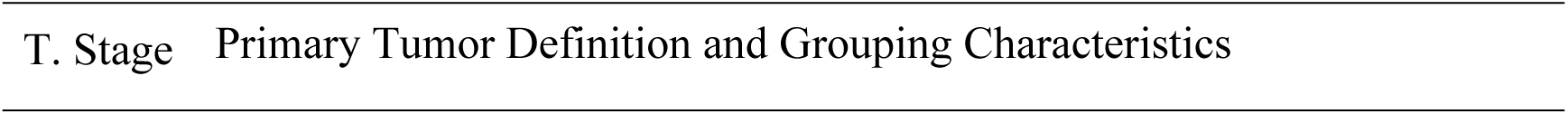
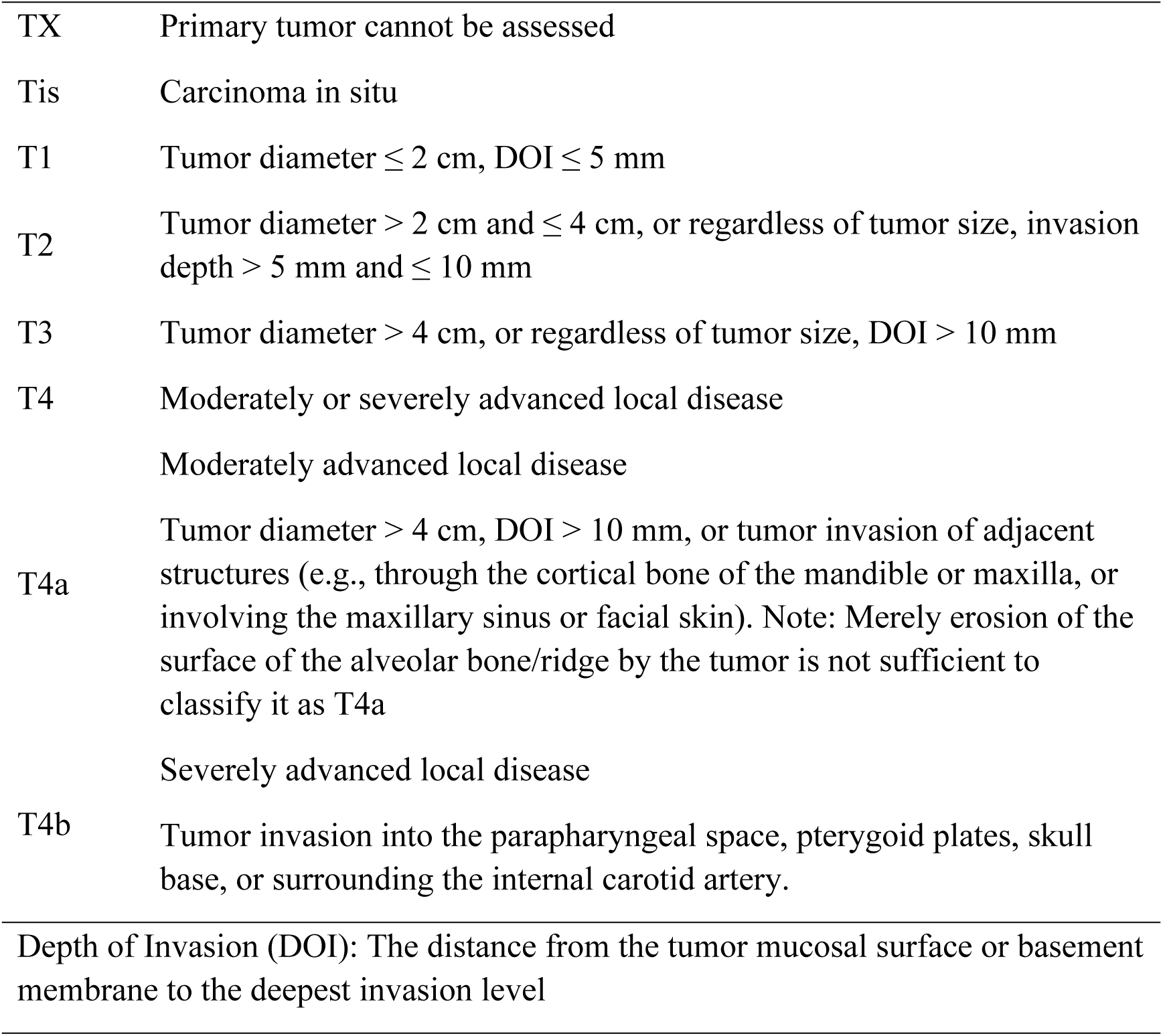
T stage definitions and characteristics in OSCC.

### Metagenomic Sequencing and Analysis

The Enzymic Universal DNAseq Library Prep Kit (KAITAI-BIO, AT4107) was utilized to process qualified samples for library construction suitable for sequencing. Initially, DNA samples underwent fragmentation, end repair, and adapter ligation, followed by PCR amplification. Target products between 300bp and 600bp were then selected and recovered using AMPure XP beads for sequencing. Post-construction, the libraries were initially quantified using Qubit, and insert fragment sizes were assessed with a Qseq100 DNA Analyzer. If the sizes aligned with expectations, the library’s effective concentration was accurately measured *via* Q-PCR to ensure quality. Upon confirming library quality, samples were mixed based on effective concentration and data requirements for downstream analysis, followed by PE150 sequencing on the NovaSeq 6000 platform. During data preprocessing, raw sequencing data were filtered using fastp software to remove low-quality sequences, yielding clean data for subsequent analysis. In the metagenomic assembly stage, clean data were assembled with megahit software, and unused reads were aligned with Bowtie2 to generate contigs. For gene prediction and abundance analysis, CDS prediction and redundancy removal were performed on contigs, and salmon software calculated abundance information for Unigenes. Species annotation was achieved by comparing Unigenes with the NR_meta database. In the functional annotation stage, DIAMOND software was employed to compare Unigenes with the GO, KEGG, and COG databases to obtain functional information. Finally, statistical and comparative analyses were conducted using methods such as PCA, Anosim, Metastats, and LEfSe to identify differences in abundance at the species and functional levels between samples.

### Cell Culture

Human tongue squamous cell carcinoma cell lines CAL27 and SCC25, human gingival fibroblast cell line HGF, and human oral keratinocyte cell line HOK were obtained from the Shanghai Institute of Biochemistry and Cell Biology and stored in liquid nitrogen. These cell lines were cultured in a constant-temperature incubator at 37°C with 95% humidity and 5% CO2. The culture medium consisted of high-glucose DMEM (Sourced, China; Gibco, USA), supplemented with 10% FBS (Bioexplore, USA) and 1% penicillin-streptomycin (Gibco, USA). After preparation, the medium was stored at 4°C.

### Bacterial Culture

*Fusobacterium nucleatum* (ATCC 25586), *Porphyromonas gingivalis* (ATCC 33277), and *Streptococcus mutans* (ATCC 25175) were sourced from the Laboratory of Oral Microecology and Systemic Diseases and preserved in liquid nitrogen. Bacterial cultures were maintained in an anaerobic chamber (Baker Ruskinn Bugbox Plus, UK) with a gas mixture of 90% N2, 5% CO2, and 5% H2. Brain Heart Infusion (BHI) solid medium was prepared by dissolving 9g in 220mL ddH2O, sterilizing under high temperature and pressure, and aliquoting into sterile 50mL centrifuge tubes for storage. Prior to experiments, bacterial cells were heat-inactivated at 75°C for 5 minutes, followed by centrifugation at 4000g for 20 minutes to collect the supernatant, designated as the heat-inactivated bacterial suspension (Hi). The supernatant was further centrifuged at 14000g for 20 minutes to obtain the bacterial culture supernatant (Supernatant). Bacterial density was measured using a spectrophotometer at OD600, with blank medium as the reference. The bacterial suspension was centrifuged at 10000 rpm for 10 minutes, the supernatant discarded, and the bacterial pellet resuspended in PBS. The resuspended pellet was transferred to 1.5 ml Eppendorf tubes, and 0.1 ml and 0.01 ml aliquots were diluted in 0.9 ml and 0.99 ml of PBS, respectively, to maintain the bacterial suspension. OD600 was then measured to determine bacterial density.

### Outer Membrane Vesicles Isolation, Observation, and Size Analysis

#### Ultracentrifugation for OMVs Isolation

*Fusobacterium nucleatum* was cultured to the third generation, from which 200 mL of bacterial suspension was processed. Sequential centrifugation steps were employed: first, cells were pelleted by centrifuging at 1000 × g for 10 minutes, and the supernatant was collected. This was followed by further clarification of the supernatant at 3000 × g for 30 minutes. The supernatant was then concentrated using a 10 kDa ultrafiltration tube, with centrifugation at 3500 rpm for 30 minutes, and vesicles were enriched by centrifuging at 10,000 × g for 30 minutes. The supernatant was subsequently filtered through a 0.22 μm pore-sized filter. Ultracentrifugation was then conducted at 100,000 × g for 1 hour at 4°C to pellet the vesicles, after which the supernatant was carefully discarded. The vesicle pellet was resuspended in PBS, and the ultracentrifugation step was repeated to ensure purity. Finally, the vesicles were resuspended in PBS, and any remaining debris was removed by low-speed centrifugation.

#### Transmission Electron Microscopy (TEM) Observation of OMVs

For TEM analysis, 10 μL of the purified outer membrane vesicles was applied to a copper grid and allowed to settle for 1 minute before the liquid was removed with filter paper. The grid was then stained with 10 μL of uranyl acetate for 1 minute, followed by another aspiration. After air-drying for several minutes, the grid was examined using a 100 kV electron microscope, and images were captured.

#### Nanoparticle Tracking Analysis (NTA)

The size distribution and uniformity of the OMVs were assessed using NTA. The frozen samples were thawed in a 25°C water bath and kept on ice. Samples were diluted in 1 × PBS and directly analyzed by NTA to determine the vesicle size distribution.

#### Cells-Bacteria Coculture

For co-culture experiments, OSCC cells (CAL27 and SCC25), human gingival fibroblasts (HGF), and human oral keratinocytes (HOK) were selected. Cells were cultured to the logarithmic phase, trypsinized to form a single-cell suspension, and then cultured for 16 to 20 hours until reaching 40% to 50% confluence. Post-digestion, cells were washed thrice with PBS, followed by further digestion with 1 mL of trypsin-EDTA until cell clumps formed, which were then dispersed *via* pipetting. Digestion was halted by adding 2 mL of complete growth medium. Cells were centrifuged, resuspended, and counted before seeding in culture plates. After 12 to 16 hours, once cells adhered, the third-generation working strains of *Fusobacterium nucleatum*, Porphyromonas gingivalis, and Streptococcus mutans were prepared into infection solutions with a multiplicity of infection (MOI) of 100, based on an optical density (OD600) of 1. The bacteria were resuspended in serum-free complete medium and added to six-well plates (1.0 mL per well), followed by incubation in a CO2 incubator at 37°C for 1 to 2 hours. Post-infection, the wells were washed 2 to 3 times with PBS, and cell culture was continued in a serum-containing complete medium.

#### Fluorescence Staining and Microscopic Observation

*Fusobacterium nucleatum* was labeled with CytoTell UltraGreen for 15 minutes at 37°C and then co-cultured with cells at an MOI of 100 for 3 to 6 hours to promote bacteria-cell interaction. Cells were subsequently fixed with 4% formaldehyde for 30 minutes to stabilize cell structure. To enhance membrane permeability, 0.1% Triton X-100 in PBS was applied on ice for 2 minutes. Nuclear staining was performed with Hoechst 33342 for 30 minutes to facilitate clear nuclear visualization under a microscope. Finally, samples were mounted and observed using a laser confocal fluorescence microscope, and images were captured with NIS-Elements software.

### Cell Proliferation Assays

#### CCK8 Assay

The CCK8 assay was utilized to assess the proliferative capacity of CAL27 and SCC25 cells. Prior to the assay, cells were cultured for 8 to 10 hours in antibiotic-free and serum-free medium to synchronize the cell cycle. After cell counting, they were seeded into a 96-well plate at a density of 3 × 10^3^ cells per well, with a volume of 100 μL per well, ensuring six replicates per group. To prevent medium evaporation, 100-150 μL of PBS was added to each well. On the third day, the medium was refreshed with a complete growth medium. Post-treatment, the medium was replaced with fresh medium containing 10% CCK-8 solution; 110 μL was added to each well, gently mixed, and incubated at 37°C for 2 hours. Absorbance was measured at 450 nm using a microplate reader, with results recorded and analyzed.

#### Plate Clone Formation Assay

The clonogenic assay evaluated the cells’ colony-forming ability. Cells were trypsinized, resuspended, and counted using a hemocytometer. A total of 200 cells were plated in 35 mm culture dishes, with three replicates per group, and 2 mL of growth medium was added to each dish. The dishes were cultured under stable conditions for two weeks, with medium changes every three days. Following 24 hours of intervention, colony formation was observed, and the culture was terminated. The dishes were washed with PBS, fixed with 1 mL of 4% paraformaldehyde for 10 minutes, and then washed three times with PBS. Giemsa staining solution was applied for 10 minutes, followed by gentle rinsing. After drying in an oven, the dishes were photographed under a microscope, and colonies were counted and analyzed, with clusters of more than 50 cells defined as colonies.

#### Cell Counting Assay

Oral squamous carcinoma cells CAL27 and SCC25 were trypsinized, resuspended, and counted before seeding at a density of 1 × 10^4^ cells per well in a 24-well plate, with 1 mL of growth medium per well. Upon cell adhesion, PBS (control group) and *Fusobacterium nucleatum* outer membrane vesicles (experimental group) were introduced into the respective wells for 24-hour co-culture. Following treatment, the supernatant was discarded, and the wells were rinsed twice with PBS. Cells were digested using 100 μL of trypsin-EDTA until complete dissociation. The resulting cell suspension was carefully pipetted to prevent bubble formation and then transferred into a hemocytometer for counting. Each sample was measured in triplicate, with the mean value used for statistical analysis.

#### RNA Sequencing

Total RNA was extracted using TRIzol reagent (Invitrogen, USA). mRNA was enriched with oligo dT magnetic beads and then fragmented into approximately 300 base pairs using a fragmentation buffer. First-strand cDNA synthesis was performed *via* reverse transcription with random primers, followed by second-strand synthesis. The resulting double-stranded cDNA underwent end repair to create blunt ends, and a single ‘A’ base was added to the 3’ end for adapter ligation. The adapter-ligated products were purified, size-selected, and subjected to PCR amplification to generate the final library. Sequencing was conducted on the Illumina NovaSeq Xplus platform. Library quantification was performed using Qubit 4.0, and clusters were generated by bridge PCR amplification on the cBot. Illumina sequencing generated a large volume of reads, which were analyzed for base distribution and quality at each cycle to assess sample sequencing and library construction quality. Raw sequencing data underwent quality control to remove adapter sequences, low-quality reads, sequences with high N content, and short sequences, resulting in high-quality clean data. These clean data were aligned with the reference genome to produce mapped data for subsequent transcript assembly and expression quantification. The alignment results were further assessed for quality, including sequencing saturation, gene coverage, and read distribution across various regions of the reference genome and chromosomes. Gene and transcript expression levels were quantified using RSEM. Differentially expressed genes were identified based on the criteria of FDR < 0.05 and |log2FC| ≥ 1, and their functions were annotated. GO and KEGG annotations were applied to these differentially expressed genes, and KEGG PATHWAY enrichment analysis was conducted using the Python scipy package.

#### Immunofluorescence Assay

CAL27 and SCC25 cells were seeded at a density of 4 × 10^4^ cells per well on 24-well plates containing cover slips. Cells were treated with *Fusobacterium nucleatum* outer membrane vesicles for 12 hours, then fixed with 4% paraformaldehyde and permeabilized with 0.2% Triton X-100. Non-specific binding sites were blocked using 10% goat serum for 30 minutes, followed by overnight incubation at 4°C with a primary anti-TNFR1 antibody diluted 1:100. After washing, cells were incubated with a secondary antibody for 1 hour. DAPI was used for nuclear staining, and the cells were observed under a microscope, with fluorescence images captured for analysis. Fluorescence intensity was quantitatively analyzed using ImageJ software.

#### TNFR1 Knockdown and Lentiviral Transduction

Specific shRNA targeting TNFR1 (sh-TNFR1) was procured from GenePharma, with the sequence tailored to precisely complement the human TNFR1 gene. A non-specific control shRNA (sh-NC) was also included. Lipofectamine 2000 was employed to co-transfect packaging plasmids with sh-TNFR1 shRNA plasmids, facilitating the construction of a lentiviral vector. CAL27 and SCC25 cells were seeded in 24-well plates at a density of 5 × 10^4^ cells per well. The lentiviral vector was then used to infect the cells at an MOI of 50:1 for 48 hours. To establish cell lines with stable TNFR1 knockdown, selection was carried out using 2 μg/mL puromycin for 3-4 weeks. The downregulation of TNFR1 mRNA and protein levels was subsequently confirmed through qPCR and Western blot analysis.

#### Western Blot

Membrane proteins were extracted using a protein extraction kit. Cells were lysed in RIPA buffer supplemented with PMSF, phosphatase inhibitors, and protease inhibitors. Pre-stained protein molecular weight markers and protein samples were loaded onto an SDS-PAGE gel, and electrophoresis was initiated at 80V, increasing to 180V once the protein markers were resolved. Following electrophoresis, proteins were transferred to a PVDF membrane. The membrane and filter paper were pre-soaked in anhydrous methanol and transfer buffer. The membrane was then incubated overnight at 4°C with a primary antibody diluted in 5% skim milk or BSA. After washing with TBST for 30 minutes, an HRP-conjugated secondary antibody, diluted in 5% skim milk, was applied for 1 hour at room temperature. Unbound secondary antibodies were washed off, and ECL reagent was applied to the membrane, followed by a 3-minute incubation at room temperature. Excess reagent was blotted off, and the membrane was pressed and preserved. The protein bands’ gray values were analyzed using ImageJ software for statistical analysis.

#### Quantitative Real-time PCR (qPCR)

For reverse transcription, genomic DNA was removed using the Takara PrimeScript RT Reagent Kit with gDNA Eraser. qPCR was performed using the Roche FastStart Universal SYBR Green Master (ROX) reagent on a Roche Light Cycler 480 II instrument. The primers were designed and synthesized by Shanghai Biotech with the following sequences (5’ to 3’):

- TNFR1 (TNFRSF1A), Forward: TCACCGCTTCAGAAAACCACC; Reverse: GGTCCACTGTGCAAGAAGAG.- GAPDH, Forward: TGCACCACCAACTGCTTAG; Reverse: GATGCAGGGATGATGTTC.

#### Animal Model and *In Vivo* Experiments

BALB/c-nude mice, aged 4 to 5 weeks, SPF grade, comprising 16 mice with an equal gender distribution, were acquired from the animal laboratory and housed in SPF facilities. The mice were randomly assigned to 4 cages, with 4 mice per cage, segregated by gender. Prior to experimentation, the mice were acclimated to the housing environment. Mice that successfully developed tumors were randomly allocated to experimental and control groups according to the following protocols: (1) Group one consisted of 3 mice each in the experimental and control groups. The experimental group received weekly injections of 50 μL of *Fusobacterium nucleatum* bacterial suspension or outer membrane vesicles with an OD600 of 1, while the control group was administered an equivalent volume of complete culture medium; (2) Group two included 5 mice each in the experimental and control groups. The experimental group received weekly injections of 50 μL of *Fusobacterium nucleatum* outer membrane vesicles, with the control group receiving an equal volume of complete culture medium; (3) Group three also consisted of 5 mice each in the experimental and control groups. The experimental group received weekly injections of 50 μL of *Fusobacterium nucleatum* outer membrane vesicles, while the control group received an equal volume of complete culture medium. The tumor cells used in this group were TNFR1-knockdown KD-CAL27 cells. Tumor dimensions were measured every 7 days, and tumor volume was calculated. Statistical analysis of the tumor volume data was performed, and the experiment was terminated once significant differences in tumor volume were observed or when the maximum tumor diameter reached 2 cm. All procedures were conducted in strict compliance with ethical and welfare standards for animal experimentation. The animal experiments adhered to a protocol approved by the NHC Key Lab of Reproduction Regulation (ethical approval number 2020-32).

#### Statistical Analysis

Statistical analysis was conducted using Prism software, version 8.4.0.671 (GraphPad Software). Depending on the data requirements, unpaired t-tests or analysis of variance (ANOVA) were employed for comparisons, with detailed descriptions provided in the figure legends. Data are expressed as mean ± standard deviation (SD). Statistical significance was indicated by * p < 0.05, ** p < 0.01, and *** p < 0.001.

### Bioinformatics Analysis

#### Data Collection

All data were obtained from public databases, including the TCGA (The Cancer Genome Atlas) HNSC dataset (TCGA-HNSC), and single-cell datasets GSE103322, GSE139324, and OSCC-GSE117257, which contain single-cell transcriptomic data for head and neck squamous cell carcinoma (HNSC) and oral squamous cell carcinoma (OSCC). The specific analysis steps and software packages used are outlined below.

#### Data Preprocessing

Gene expression data were normalized using the scale function in R to convert the expression levels into z-scores, standardizing the data across different datasets. The normalization was performed using the formula (x - \mu) / \sigma(x−μ)/σ, where xx is the individual data point, \muμ is the mean, and \sigmaσ is the standard deviation. Z-scores were used for subsequent differential expression and pathway enrichment analyses.

#### Differential Expression Analysis

Differential expression analysis was performed using the limma package in R to calculate the log2 fold change (log2FC) between different sample groups. Genes with |log2FC| > 0.5 and an adjusted p-value (FDR) < 0.05 were considered differentially expressed. The significance of gene expression differences was evaluated using the Wilcoxon rank-sum test.

#### Kaplan-Meier Survival Analysis

Kaplan-Meier survival analysis was performed using the survival and survminer packages in R to evaluate the association between TNFRSF1A (TNFR1) expression and patient outcomes, including overall survival (OS), disease-specific survival (DSS), and progression-free interval (PFI) in head and neck cancer. Univariate Cox regression analysis was conducted to calculate hazard ratios (HR) and 95% confidence intervals (CI). Log-rank tests were used to assess the significance of survival differences, with p-values < 0.05 considered statistically significant.

#### Correlation Analysis

Correlation analysis was performed using the cor.test() function in R. Pearson’s correlation test was used for normally distributed data, while Spearman’s correlation was applied for non-normally distributed data or data with different units of measurement. Comparisons between discrete variables (e.g., tumor vs. normal) were conducted using the non-parametric Wilcoxon rank-sum test.

#### Gene Set Enrichment Analysis (GSEA) and Gene Set Variation Analysis (GSVA)

Gene set enrichment analysis (GSEA) was performed using the clusterProfiler package in R, with enrichGO and enrichKEGG functions used to assess Gene Ontology (GO) and Kyoto Encyclopedia of Genes and Genomes (KEGG) pathway enrichment. The GSVA package was used to evaluate the activity of cancer-related pathways across multiple sample datasets. All pathway activity scores were standardized using z-scores.

#### Single-Cell RNA Sequencing (scRNA-seq) Data Analysis

Single-cell transcriptomic data were processed and analyzed using the Seurat package in R. Uniform Manifold Approximation and Projection (UMAP) dimensionality reduction was performed using the RunUMAP function to project high-dimensional data into a two-dimensional space, enabling the identification of distinct cell populations. Differential expression analysis of cell types was used to identify marker genes (e.g., MKI67). Each point on the UMAP plot represents a single cell, with colors indicating gene expression levels, where red represents high expression and gray represents low expression. Cell populations were identified using the FindClusters function.

#### Intercellular Communication Analysis

Intercellular communication analysis was performed using the CellChat package, which analyzes the signaling networks between cells based on a receptor-ligand interaction database. Ligand-receptor interaction networks were constructed to further evaluate the signaling mechanisms of TNFRSF1A and its transmission across different cell types.

#### Statistical Analysis

All statistical analyses were performed in R using two-sided tests. P-values less than 0.05 were considered statistically significant, with p-values less than 0.01, 0.001, and 0.0001 indicating high significance. Statistical results were reported as *P < 0.05, **P < 0.01, ***P < 0.001, and ****P < 0.0001. All data analysis was conducted using R (version 4.3.3). Data processing and visualization were performed using the ggplot2, Seurat, limma, clusterProfiler, survival, survminer, GSVA, and CellChat R packages.

## Results

### Dysbiosis of Oral Microbiota and Functional Alterations in OSCC Progression

During the progression of OSCC, significant alterations in the oral microbiota structure occur, which may have profound implications for tumor development. A comprehensive analysis of the microbial community structure in samples from healthy normal tissues (NT) and various OSCC stages (T1-T4) was conducted. Notably, *Fusobacterium nucleatum* (*F. nucleatum*), *Streptococcus* mitis (*S. mitis*), and *Escherichia coli* (*E. coli*) exhibited a marked increase in relative abundance as OSCC progressed (Figure 1.A). Among these, *F. nucleatum* showed a particularly pronounced increase compared to normal tissue across different cancer stages. PCA and ANOSIM further corroborated the significant differences in the oral microbial community structure (Figure 1.B, C). Metastats analysis revealed significant variations in the relative abundance of *F. nucleatum* across OSCC stages, especially in the early stages (Figure 1.E). LEfSe discriminant analysis highlighted substantial differences in microorganisms, such as *F. nucleatum*, among the different stages (Figure 1.D).

**Figure 1.**
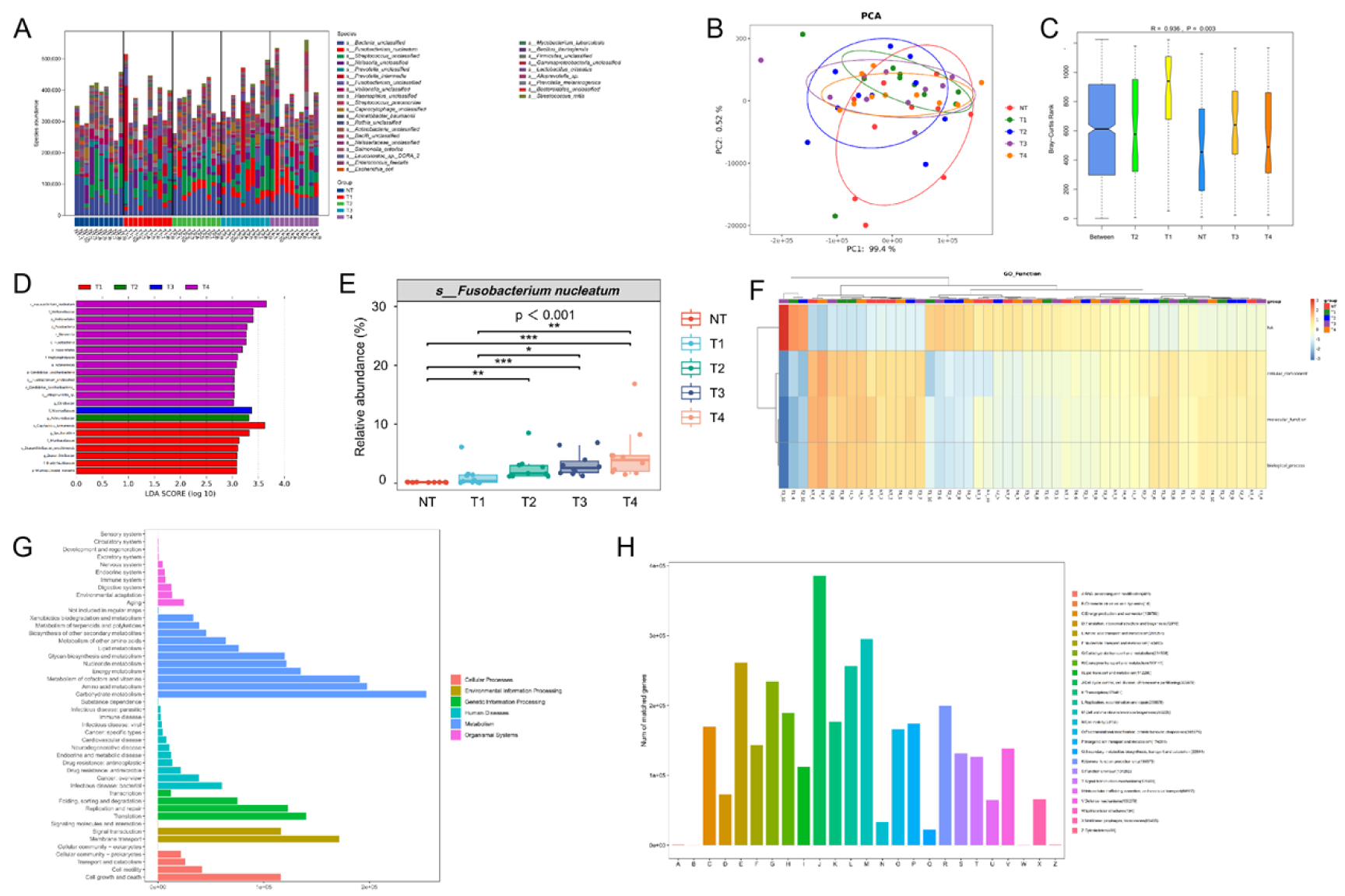
Overview of Oral Microbiota Structure and Functional Dynamics in OSCC Progression. A. Stacked bar chart illustrating the relative abundance of microbial species across different stages of OSCC, highlighting changes in oral microbiota community structure; B. Principal Component Analysis (PCA) plot comparing OSCC stages with normal tissue samples, showcasing significant β-diversity differences among the groups; C. ANOSIM analysis boxplot based on species abundance, demonstrating microbial community structure differences between normal tissue (NT) and various OSCC stages (T1-T4); D. Boxplot depicting the variation in relative abundance of *Fusobacterium nucleatum* across OSCC stages, analyzed using the Metastats method; E. Bar plot of LDA analysis revealing significant differences in microbial taxa between different OSCC stages; F. Heatmap of GO functional classification, reflecting functional expression patterns of the oral microbiota in various biological processes during OSCC progression; G. Distribution of oral microbiota community functions at the KEGG pathway level, highlighting key biological processes and pathways involved in OSCC development; H. Bar chart of COG functional classification, detailing the distribution of Unigene counts across different functional categories within the oral microbiota community. (*P < 0.05, **P < 0.01, ***P < 0.001).

Functional annotation analysis using the GO, KEGG, and COG databases identified characteristic functional changes in the oral microbiota between normal tissue (NT) and different OSCC stages (T1-T4). GO heatmap analysis illustrated distinct expression patterns of specific GO functional categories across various cancer stages, suggesting that certain functions may be pivotal in cancer progression (Figure 1.F). KEGG pathway annotation results indicated extensive unigene involvement in cellular processes and metabolic pathways, underscoring the relevance of these biological processes in OSCC development (Figure 1.G). COG functional classification analysis further revealed significant alterations in the number of unigenes associated with protein biosynthesis, carbohydrate metabolism, transcription, and other processes, providing a foundation for the in-depth exploration of the molecular mechanisms underlying OSCC (Figure 1.H).

### Influence of Oral Microbiota on OSCC Cell Proliferation

Based on microbiome sequencing of clinical samples, *Fusobacterium nucleatum* (*Fn*), *Porphyromonas gingivalis* (*Pg*), and *Streptococcus mutans* (*Sm*) were co-cultured with OSCC cells to evaluate their impact on OSCC proliferation (Figure 2A). The CCK8 assay results demonstrated that, compared to the control group, treatment with *Fn*, *Fn culture* supernatant (Suprnt), and heat-inactivated *Fusobacterium nucleatum* (*hiFn*) significantly enhanced the proliferation of CAL27 and SCC25 cells (Figure 2B). Specifically, the Fn treatment group markedly promoted proliferation in both cell lines; similarly, the Suprnt and hiFn treatment groups exhibited significant proliferative effects on both cell lines. The Pg treatment group significantly promoted proliferation in CAL27 cells but did not have a notable effect on SCC25 cells, while the Sm treatment group only slightly increased proliferation in CAL27 cells.

**Figure 2.**
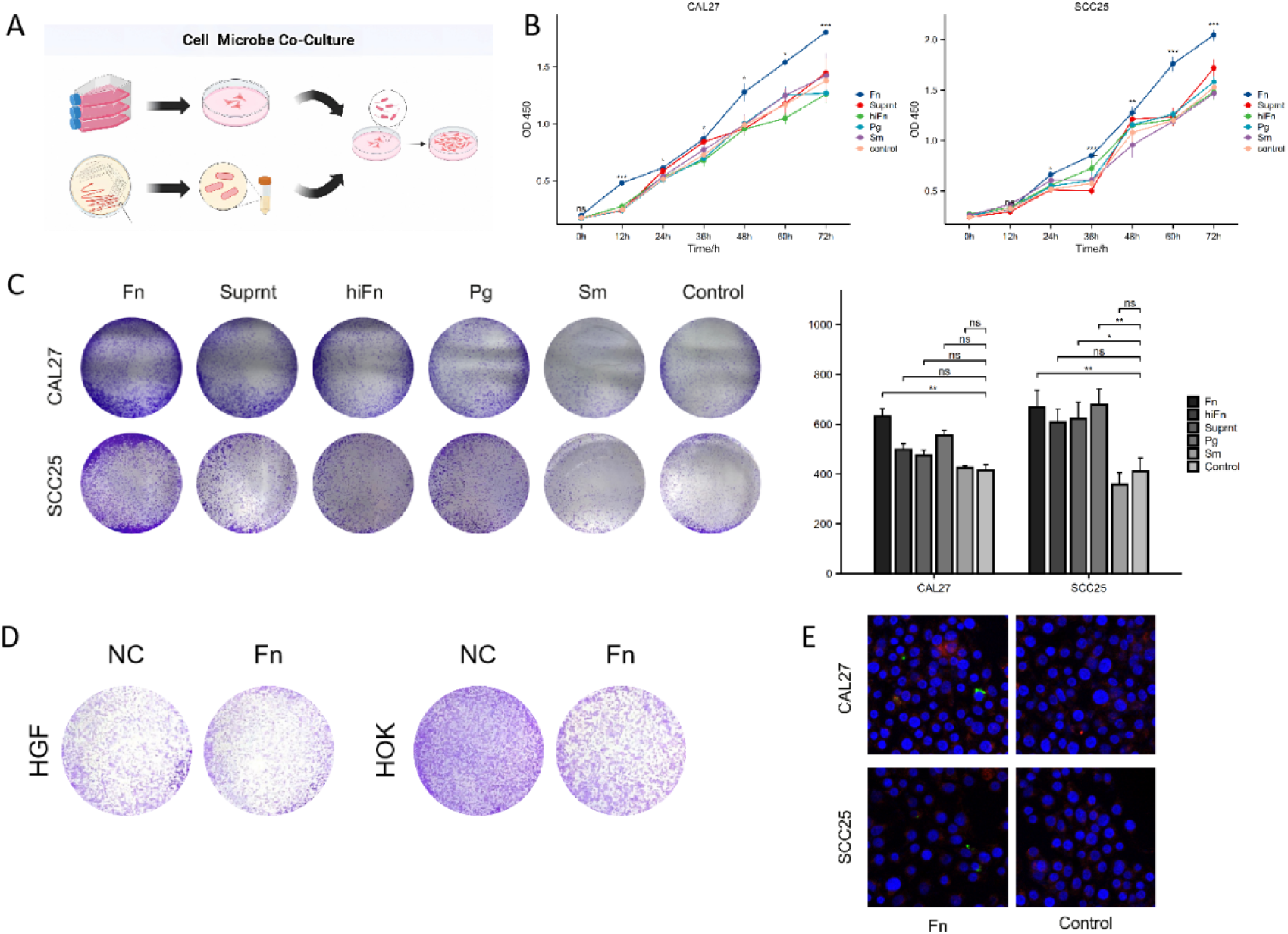
Changes in Proliferative Capacity of CAL27 and SCC25 Cells under Microbial Intervention. A. Schematic diagram illustrating the co-culture setup of CAL27 and SCC25 cells with various microorganisms; B. CCK8 proliferation curves depicting the time-dependent growth of CAL27 and SCC25 cells under different treatment conditions, including Fn, Suprnt, hiFn, Pg, Sm, and control; C. Colony formation efficiency plot showing the colony-forming ability of CAL27 and SCC25 cells across various treatments; D. Comparison of the proliferative capacity of normal oral cell lines HGF and HOK under Fn treatment conditions; E. Fluorescence staining images displaying nuclear staining of CAL27 and SCC25 cells post-treatment with Fn and control, indicating that Fn did not internalize but exerted its effects by adhering to the cell surface.

In the colony formation assay, both Fn and Suprnt treatments significantly boosted the colony-forming ability of CAL27 cells (Figure 2C), whereas the hiFn, Pg, and Sm treatment groups did not show significant effects in CAL27 cells. In SCC25 cells, Fn and Suprnt treatments also significantly enhanced colony formation compared to the control group (Figure 2C). Importantly, Fn did not exhibit a significant proliferative effect on normal oral cell lines HGF and HOK (Figure 2D).

Fluorescence staining indicated a marked increase in the number of cell nuclei in CAL27 and SCC25 cells following Fn treatment, suggesting an elevated proliferation rate, while the control group showed relatively fewer cell nuclei (Figure 2E). Additionally, observations revealed that Fn did not directly enter the cells; the green-stained *Fusobacterium nucleatum* was found attached to the cell surface, implying that its proliferation-promoting effect may occur through indirect mechanisms rather than direct cell entry. This observation is further supported by the proliferation-promoting effects observed with the culture supernatant (Suprnt), suggesting that *Fusobacterium nucleatum* may secrete certain substances that exert their effects through the supernatant (Figure 2E).

### Fn-OMVs Preparation and *In Vivo* and *In Vitro* Intervention in OSCC

The effects of OMVs on OSCC cells were rigorously assessed using transmission electron microscopy, nanoparticle size analysis, colony formation assays, CCK-8 proliferation curves, and *in vivo* tumor formation experiments in nude mice. Transmission electron microscopy revealed that OMVs possess distinct double-membrane structures and varying vesicle morphologies, highlighting their complexity and biosynthetic diversity (Figure 3A). Nanoparticle size analysis demonstrated consistent stability at specific concentrations, with an average diameter of 156.7 ± 7.15 nanometers, supporting further biological applications (Figure 3B). Colony formation assays showed that cells treated with Fn and its OMVs exhibited varying capacities for colony formation, with OMVs displaying a slightly stronger proliferation-promoting effect than Fn (Figure 3C). CCK-8 proliferation curves further confirmed significant differences in cell proliferation rates compared to controls, underscoring OMVs’ more pronounced regulatory impact on cell proliferation than Fn (Figure 3D).

**Figure 3.**
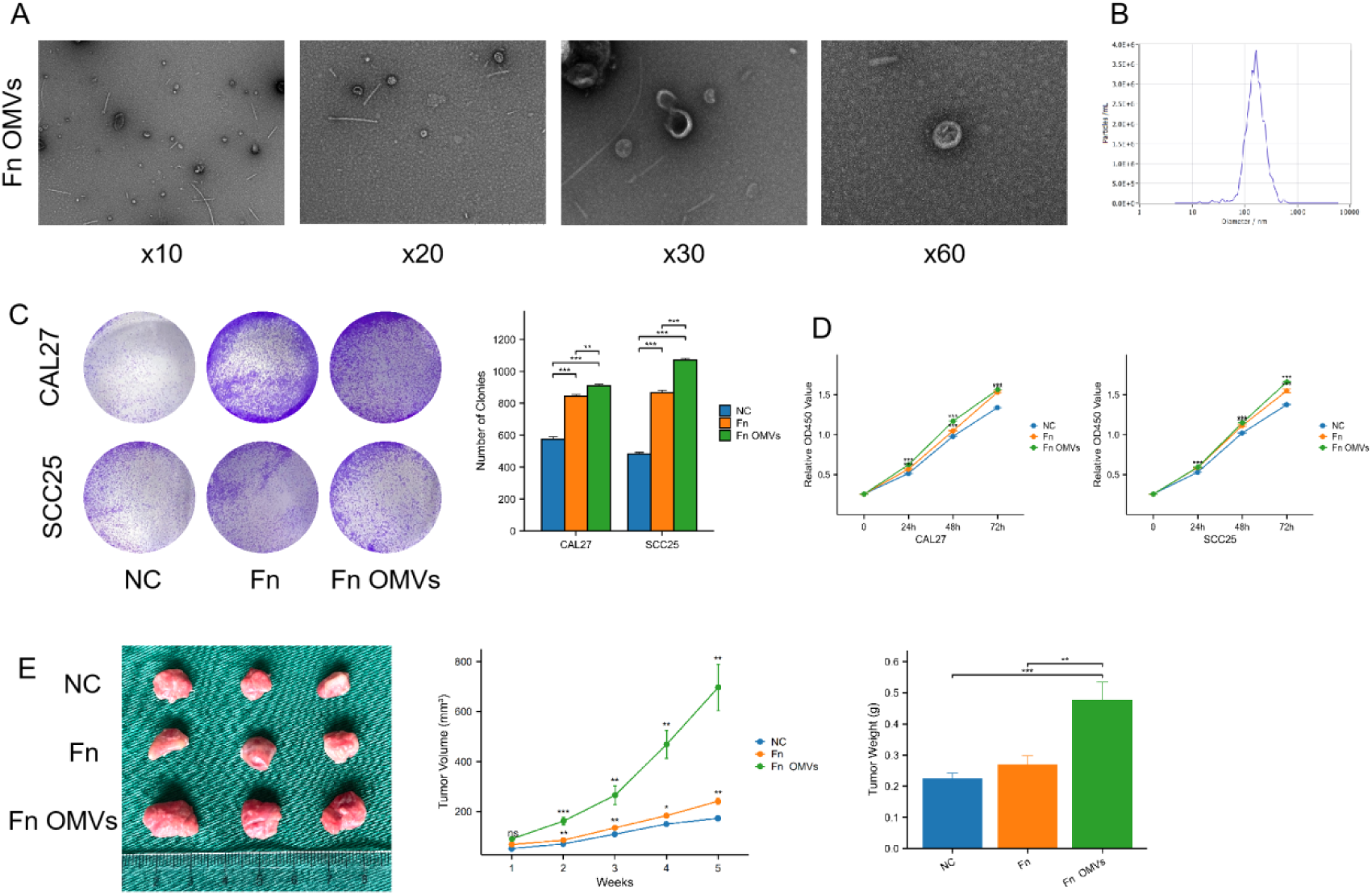
Effects of Fn OMVs on Cell Proliferation Behavior in *In Vivo* and *In Vitro* Experiments. A. Morphological characteristics of outer membrane vesicles (OMVs) observed under transmission electron microscopy (Voltage = 80kV, magnification = × 10k, × 20k, × 30k, × 60k); B. Histogram depicting nanoparticle size distribution along with representative transmission electron microscopy images. The average nanoparticle diameter is 156.7 (± 7.15) nanometers; C. Colony formation assay results showing the number of cell clones after treatment with *Fusobacterium nucleatum* and its OMVs, indicating an enhanced colony formation capacity post-treatment; D. CCK-8 cell proliferation curve demonstrating that cells treated with OMVs exhibit accelerated proliferation compared to the control group; E. *In vivo* tumor formation results in nude mice, showing tumor volumes within 5 weeks post-injection. Tumors in the treated groups are significantly larger than those in the control group, indicating that *Fusobacterium nucleatum* and its OMVs significantly promote tumor growth.

*In vivo* subcutaneous tumor formation experiments in nude mice validated these results, with both Fn-treated and Fn OMVs-treated groups exhibiting significantly larger tumor volumes than controls. The OMVs-treated group displayed particularly pronounced differences, suggesting a potent tumor-promoting effect of OMVs (Figure 3E).

### FnOMVs Induced OSCC Proliferation: RNA-Seq Analysis and Validation of TNFR1 by RT-qPCR and Western Blot

Transcriptional profiling of CAL27 cells treated with OMVs revealed significant alterations in gene expression, as depicted in the heatmap of sequenced genes (Figure 4.A). Notable changes were observed in tumor-related genes such as TNFR1 and WNT9A, along with several non-coding RNAs and genes linked to cancer progression. Volcano plots highlighted the upregulation of genes involved in cell proliferation, apoptosis, and signaling pathway regulation (Figure 4.B). RT-qPCR confirmed a significant increase in TNFR1 mRNA expression following OMVs treatment in both CAL27 and SCC25 cell lines (Figure 4.C). Western blot analysis further validated these outcomes, demonstrating elevated TNFR1 protein levels in the OSCC cell lines CAL27 and SCC25 post-OMVs treatment (Figure 4.D).

**Figure 4.**
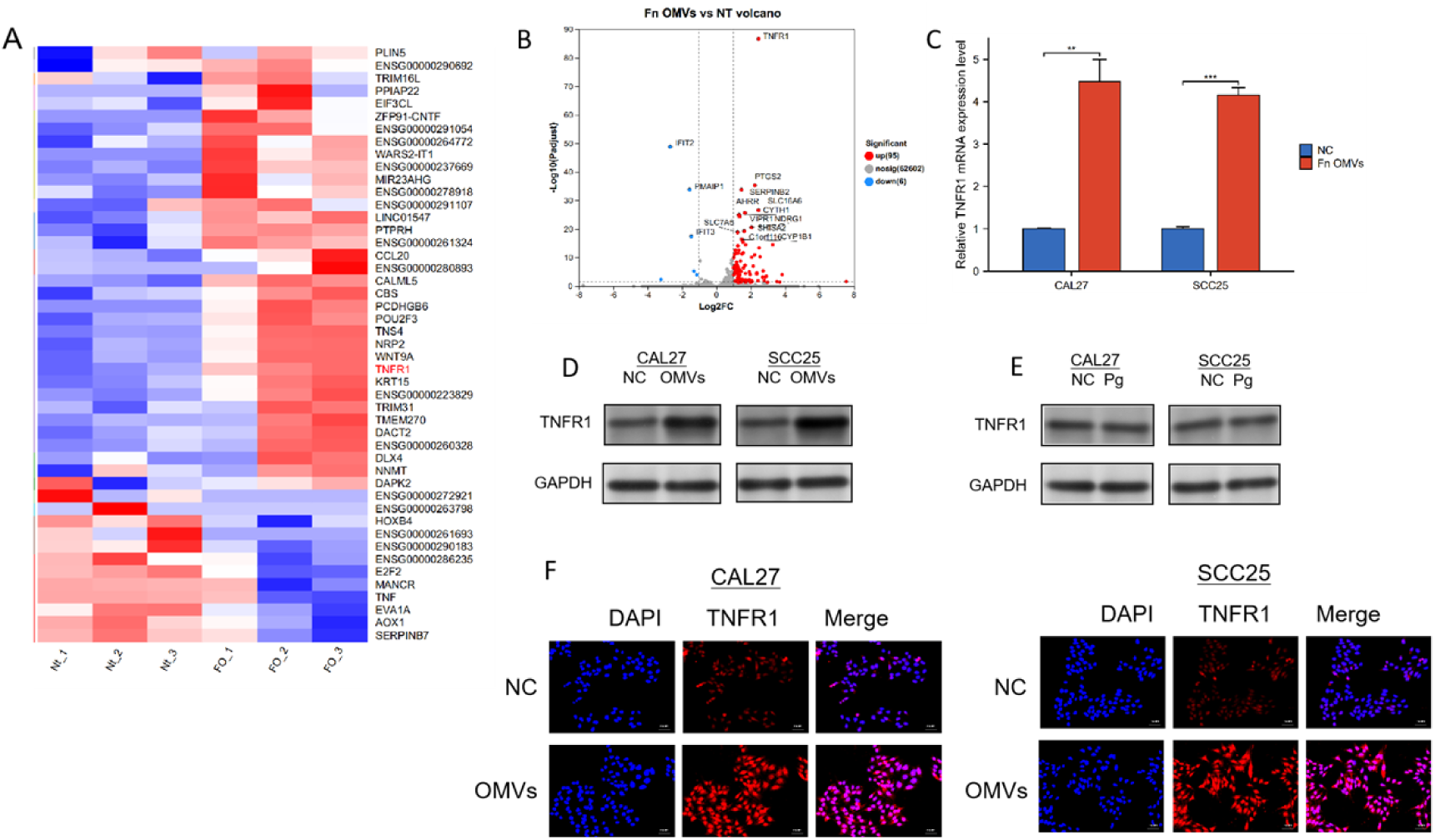
Analysis of Gene and Protein Expression Levels Regulated by Outer Membrane Vesicles in OSCC Cells. A. Differential gene expression levels between the control group (NC) and the OMVs-treated group. Red indicates upregulated genes, while blue denotes downregulated genes; B. Volcano plot illustrating changes in gene expression following OMVs intervention. Significantly upregulated genes are positioned in the upper right corner, and significantly downregulated genes are in the upper left corner; C. TNFR1 mRNA expression levels in CAL27 and SCC25 cell lines after OMVs treatment, showing significant differences between the OMVs-treated group and the control group; D. Western blot analysis depicting the differential expression of TNFR1 protein in CAL27 and SCC25 cell lines between the OMVs-treated group and the control group; E. TNFR1 protein expression levels in CAL27 and SCC25 cell lines under conditions with and without Porphyromonas gingivalis (Pg); F. Immunofluorescence staining revealing the spatial distribution of TNFR1 protein in CAL27 and SCC25 cell lines. DAPI staining indicates cell nuclei (blue), TNFR1 is labeled in red, and the merged image shows their overlap.

To assess cellular responses under different conditions, Porphyromonas gingivalis (Pg) was used to intervene in OSCC cells, specifically evaluating its impact on TNFR1 protein expression. The results indicated that Pg did not significantly alter TNFR1 protein levels under the experimental conditions (Figure 4.E).

Immunofluorescence staining provided insights into the spatial distribution of TNFR1 within the cells. Cells treated with OMVs exhibited a stronger TNFR1 signal, particularly in regions near the cell nucleus (Figure 4.F). This suggests that OMV treatment may enhance the aggregation or surface expression of TNFR1, aligning with its role in cell signaling.

### OMVs Regulate OSCC Proliferation *via* TNFR1

Lentiviral transfection was utilized to knock down the TNFR1 gene in OSCC, and the expression of TNFR1 protein in CAL27 and SCC25 oral squamous cell carcinoma cell lines was assessed *via* Western blotting. The results demonstrated a significant reduction in TNFR1 protein levels in the TNFR1 knockdown group (KD), while levels remained elevated in the control group (NC), confirming the efficacy of the knockdown (Figure 5A). Cell counting experiments were conducted to compare the proliferation rates between the control group (NC), the knockdown group (KD), and the knockdown group post-treatment with outer membrane vesicles (OMVs). The knockdown group (KD) exhibited a marked reduction in the proliferation ability of both CAL27 and SCC25 cell lines. However, the addition of OMVs significantly enhanced the proliferation in the knockdown group, particularly at 48 and 72 hours (Figure 5B).

**Figure 5.**
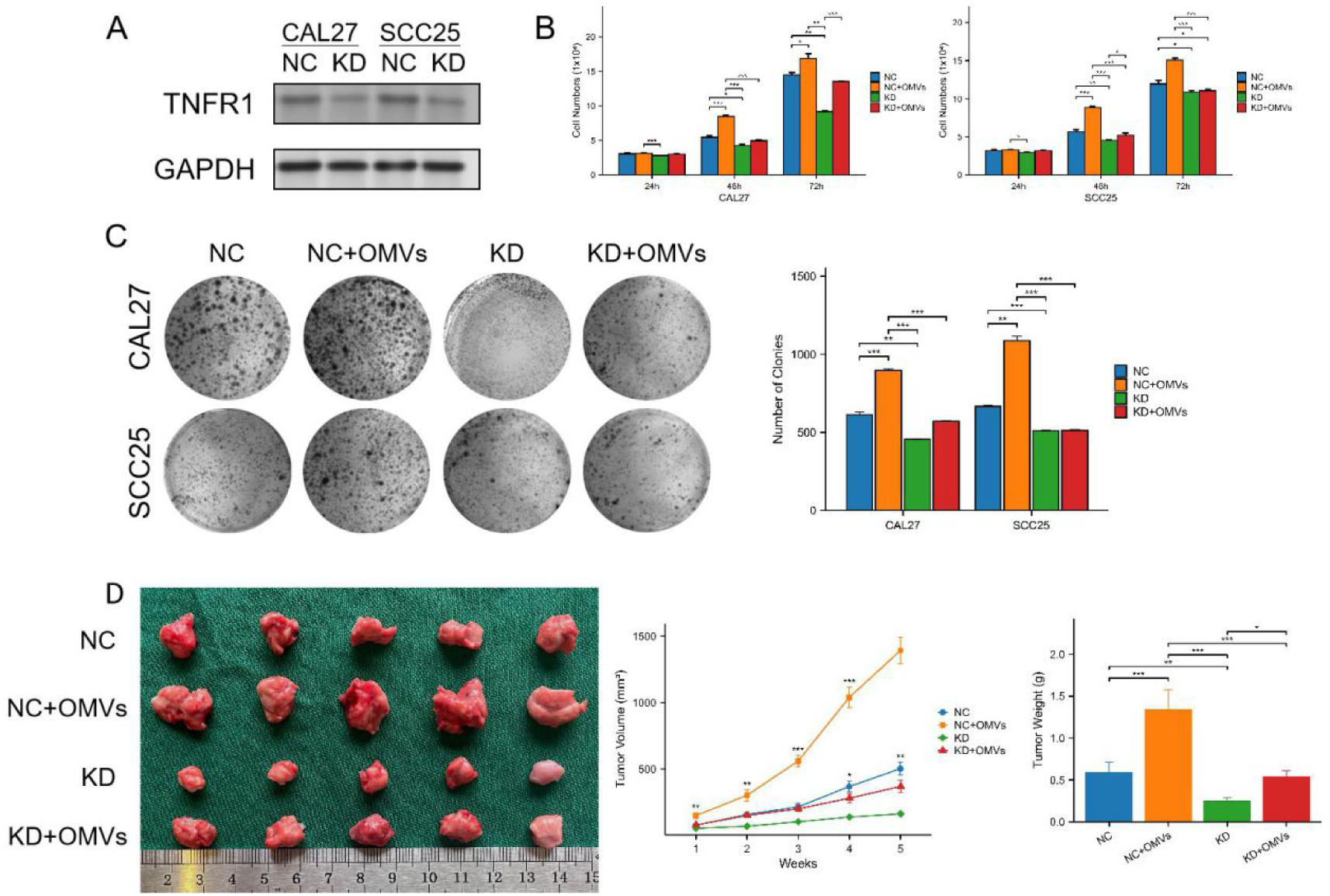
Regulation of Proliferation and Tumor Formation in Oral Squamous Cell Carcinoma Cells by Outer Membrane Vesicles *via* TNFR1. A. Western Blot analysis of TNFR1 protein levels comparing the control group (NC) with the TNFR1-knockdown group (KD), using GAPDH as an internal reference to ensure consistent protein loading; B. Cell counting analysis of CAL27 and SCC25 cells at 24, 48, and 72 hours, showing that TNFR1-knockdown cells (KD) exhibited a significantly increased proliferation ability compared to the control group (NC) after treatment with outer membrane vesicles (OMVs), particularly at 48 and 72 hours; C. Cloning formation ability analysis, indicating a higher number of clones in TNFR1-knockdown cells (KD) compared to the control group (NC) following treatment with OMVs; D. Comparison of tumor volume trends in subcutaneous tumor formation experiments in nude mice among the control group (NC), outer membrane vesicle-treated group (NC+OMVs), knockdown group (KD), and knockdown group with subsequent outer membrane vesicle treatment (KD+OMVs).

Furthermore, the number of colonies in CAL27 and SCC25 cell lines significantly decreased in the knockdown group (KD). Conversely, treatment with OMVs substantially increased colony formation in the knockdown group, bringing the number of colonies close to or even surpassing that of the control group (Figure 5C).

Subcutaneous tumor formation experiments in nude mice revealed a slower tumor growth rate in the knockdown group (KD). However, OMV treatment significantly accelerated tumor growth, with the most rapid tumor progression observed in the knockdown group treated with OMVs (KD + OMVs) (Figure 5D).

### Fn OMVs regulate the NF-κB signaling pathway to promote OSCC proliferation

KEGG pathway enrichment analysis of the non-knockdown OSCC transcriptome at various intervention times revealed the activation of several key signaling pathways, including NF-κB, AKT/mTOR, MAPK/ERK, and JNK. The activation of these pathways not only influences cell inflammation and survival but may also be linked to processes such as cell proliferation and apoptosis, providing preliminary evidence for how outer membrane vesicles (OMVs) regulate cell fate through multiple signaling pathways. Particular focus was given to the NF-κB signaling pathway, which exhibited the most significant changes following OMVs intervention. By examining different time points, the study uncovered the dynamic interplay between OMVs and the NF-κB signaling pathway (Figure 6A). The pathway’s activation status fluctuated during the 3 to 24-hour intervention period, indicating that OMVs impact both short-term and long-term activation, with the pathway remaining continuously activated throughout the intervention (Figure 6A).

**Figure 6.**
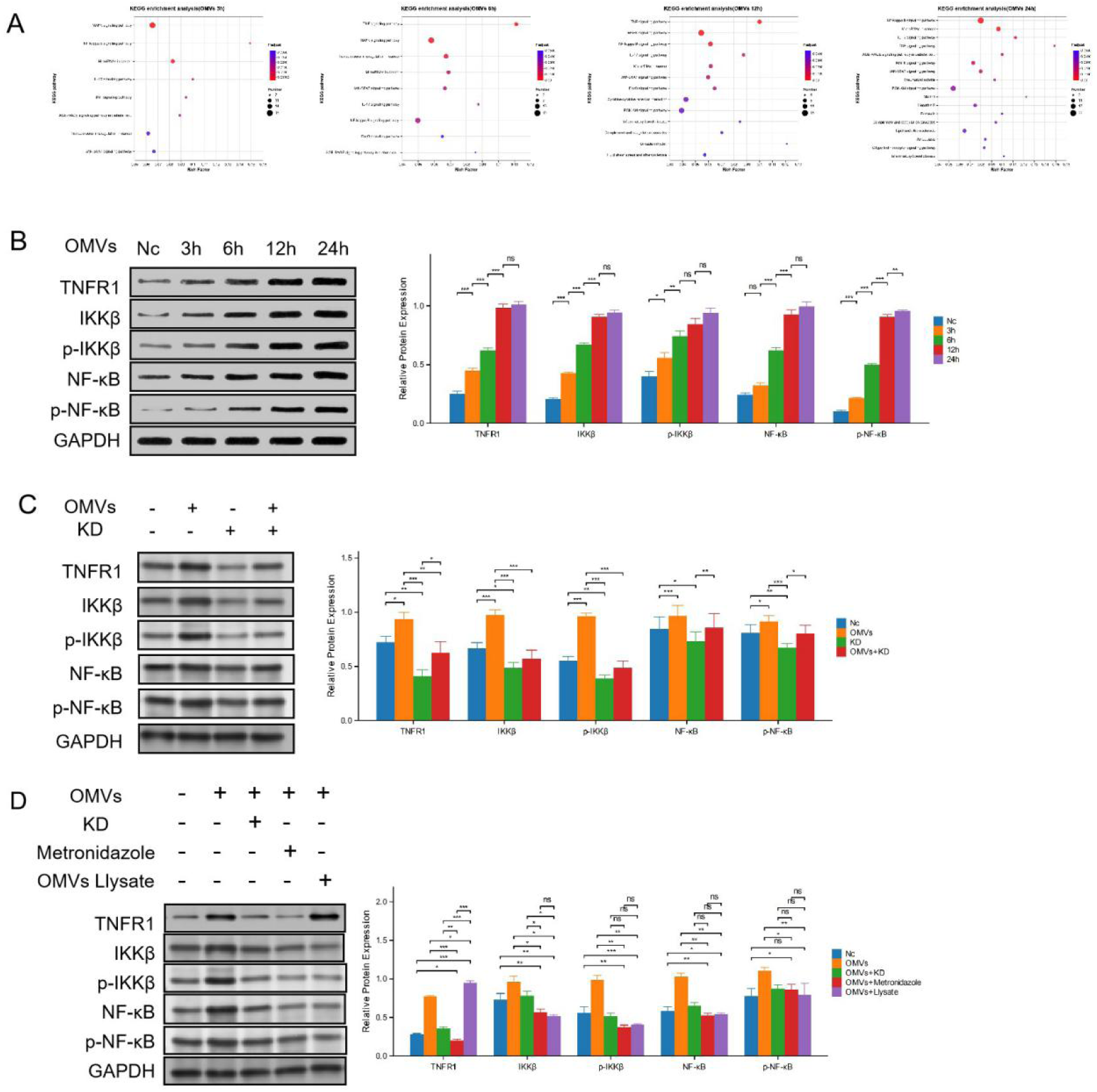
Outer Membrane Vesicles (OMVs) Promote the Proliferation of OSCC cells (CAL27) by Regulating the NF-κB Signaling Pathway. A. KEGG pathway enrichment results in the transcriptome following OMVs intervention, highlighting key pathways affected; B. Bar graph displaying changes in the expression levels of TNFR1, IKKβ, p-IKKβ, NF-κB, and p-NF-κB under OMVs treatment, comparing untreated cells with those treated at various time points; C. Bar graph illustrating the impact of TNFR1 knockout (KD) on OMVs-induced activation of the NF-κB signaling pathway, comparing untreated cells with those treated with OMVs; D. Bar graph detailing the effects of OMVs, TNFR1 gene knockout, metronidazole treatment, and OMVs protein protease on the NF-κB signaling pathway, showcasing the combined effects of different treatments on pathway activity.

Further analysis of key proteins within the NF-κB signaling pathway was conducted at various time points (no intervention, 3 hours, 6 hours, 12 hours, and 24 hours) (Figure 6B). TNFR1 (tumor necrosis factor receptor 1) expression remained relatively unchanged over time. However, the expression of IKKβ (IκB kinase β) and its phosphorylated form, p-IKKβ, increased notably after OMVs treatment, particularly at 12 and 24 hours. While the total NF-κB levels did not significantly change, its phosphorylated form, p-NF-κB, exhibited an increasing trend over time, especially at 12 and 24 hours, signifying the activation of the NF-κB signaling pathway.

Subsequent experiments validated the changes in the NF-κB signaling pathway following TNFR1 gene knockout. The knockout of TNFR1 significantly diminished the activation of the NF-κB pathway induced by OMVs, underscoring TNFR1’s pivotal role in OMV-mediated signaling pathway activation (Figure 6C). While TNFR1 expression slightly increased following OMVs treatment, it remained unchanged under knockout conditions. The increase in IKKβ and p-IKKβ expression observed after OMVs treatment was inhibited under KD conditions. Similarly, the elevated expression of NF-κB and p-NF-κB following OMVs treatment was either inhibited or reduced in the TNFR1 knockout context.

To further explore alternative methods for inhibiting OSCC proliferation, the study examined the effects of OMVs intervention, TNFR1 gene knockout, metronidazole (an effective antibacterial drug against *Fusobacterium nucleatum*), and bacterial lysate (degraded OMVs protein) on the NF-κB signaling pathway (Figure 6D). The findings revealed that in the presence of OMVs, both metronidazole and bacterial lysate significantly inhibited NF-κB pathway activation. Similarly, TNFR1 gene knockout exhibited a substantial inhibitory effect on this pathway. OMVs treatment led to an increase in TNFR1 and IKKβ expression, but this increase was inhibited by both gene knockout and metronidazole treatment. OMVs lysate treatment resulted in elevated TNFR1 expression, with a lesser effect on IKKβ levels. The expression pattern of p-IKKβ mirrored that of IKKβ, but the increase was more pronounced following OMVs lysate treatment. NF-κB expression increased after OMVs treatment but decreased following TNFR1 knockout and metronidazole treatment, while the effect of OMVs lysate on NF-κB was minimal. Expression of p-NF-κB significantly increased after OMVs treatment but was reduced following gene knockout and metronidazole treatment. OMVs lysate also caused an increase in p-NF-κB expression.

These results suggest that OMVs from *Fusobacterium nucleatum* regulate the NF-κB signaling pathway by activating specific signaling proteins and that specific gene knockdown or drug treatment can modulate this regulation. Such changes may influence cellular processes related to inflammation, proliferation, and survival.

### Expression of TNFR1 and Its Clinical Significance in HNSCC

TNFRSF1A (TNFR1) is significantly upregulated in head and neck squamous cell carcinoma (HNSC). Differential expression analysis of the TCGA-HNSC dataset reveals that TNFRSF1A expression is significantly higher in tumor tissues compared to normal tissues (Wilcoxon test, p < 0.0001) (Figure 7A-B). Paired sample analysis further validates the consistent upregulation of this gene in tumor samples from patients. Immune microenvironment component distribution analysis reveals that high TNFRSF1A expression is closely associated with an increased proportion of immune-related cells, particularly monocyte-macrophages and fibroblasts (Figure 7C). Kaplan-Meier survival analysis indicates that high TNFRSF1A expression is significantly correlated with worse overall survival (OS), disease-specific survival (DSS), and progression-free interval (PFI) (p < 0.01) (Figure 7D). Analysis across multiple single-cell datasets (HNSC_GSE103322, GSE139324, and OSCC_GSE117257) reveals that TNFRSF1A is predominantly expressed in malignant cells and the monocyte-macrophage subpopulations (Figure 7E–H). Gene set variation analysis (GSVA) suggests that pathways related to inflammatory responses and TNF signaling are significantly activated in samples with high TNFRSF1A expression (Figure 7F). These findings underscore the pivotal role of TNFRSF1A in tumor immune microenvironment remodeling and prognosis in HNSCC.

**Figure 7.**
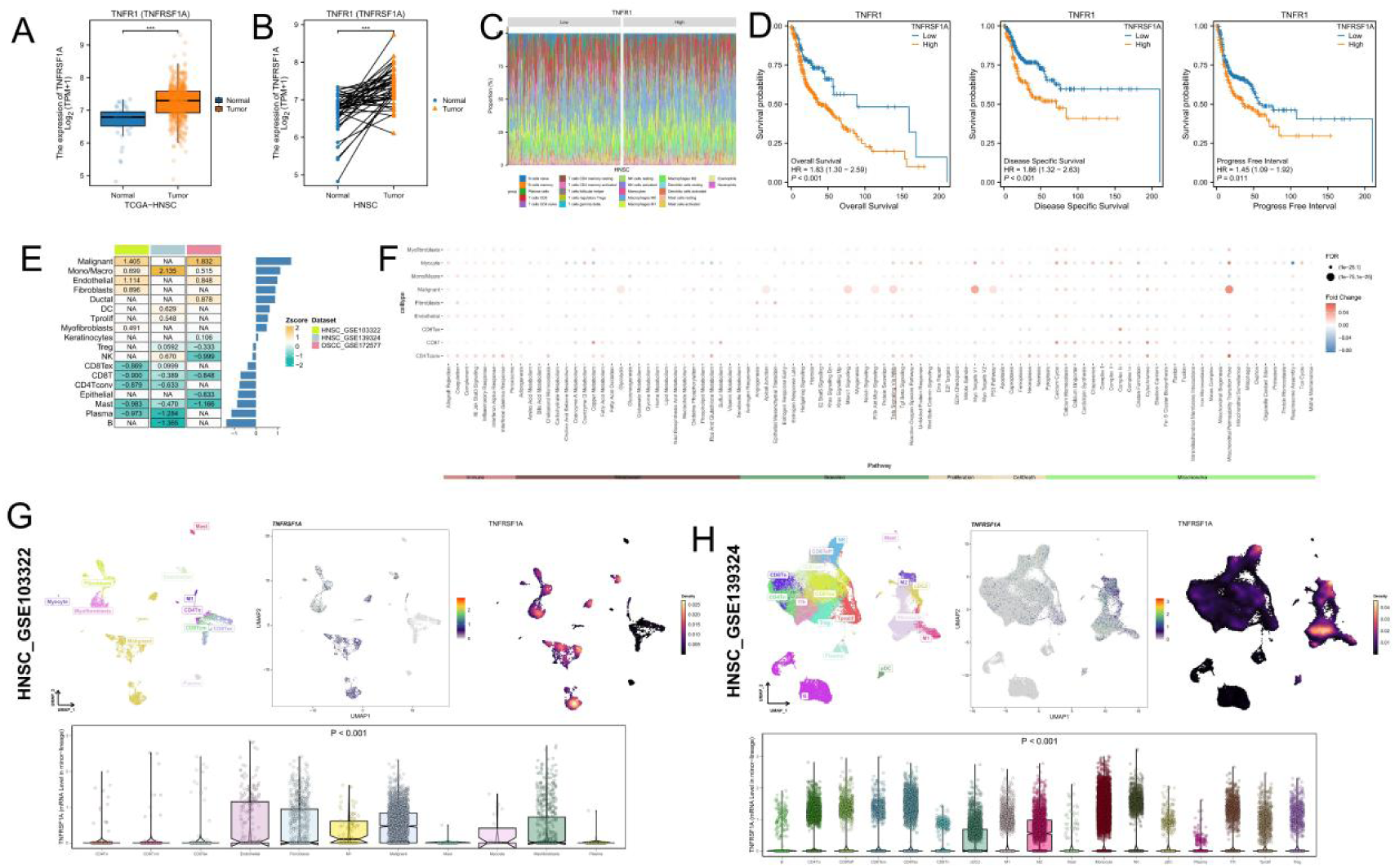
TNFRSF1A expression and prognostic significance in head and neck squamous carcinoma (HNSC). (A–B) Boxplots showing TNFRSF1A expression levels in normal and tumor tissues from TCGA-HNSC cohort and paired samples. (C) Immune cell infiltration profiles stratified by TNFRSF1A expression (high vs low). (D) Kaplan–Meier survival curves for overall survival (OS), disease-specific survival (DSS), and progression-free interval (PFI) in high vs low TNFRSF1A expression groups. (E–H) Single-cell expression profiles of TNFRSF1A in HNSC datasets (GSE103322 and GSE139324). UMAP visualization shows TNFRSF1A distribution across different cell types; violin plots indicate quantitative expression differences. (F) Pathway enrichment analysis (GSVA) of TNFRSF1A-associated signatures. *P < 0.05, **P < 0.01, ***P < 0.001, ****P < 0.0001.

### Expression of TNFR1/TNF Signaling in OSCC

At the single-cell level in oral squamous cell carcinoma (OSCC), the expression and signaling patterns of TNFRSF1A across different cell subpopulations were further explored. UMAP dimensionality reduction results show that TNFRSF1A is highly expressed in malignant cells and certain immune cell populations, particularly M1 macrophages and CD8+ T cells (Figure 8A). Intercellular communication network analysis reveals significant ligand-receptor interactions between TNFRSF1A+ malignant cells and other cell types such as monocyte-macrophages and fibroblasts (Figure 8B–C). TNFRSF1A+ cells also exhibit high outgoing signaling intensity and centrality within the TNF signaling network (Figure 8D–G). Further signaling pathway analysis indicates robust bidirectional signaling activity of TNFRSF1A-related pathways in both malignant and immune cells (Figure 8E–F). In the TNF signaling network, TNFRSF1A+ malignant cells are identified as key signal senders and mediators (Figure 8I–J), suggesting that TNFRSF1A may promote tumor progression by regulating tumor-immune cell interactions.

**Figure 8.**
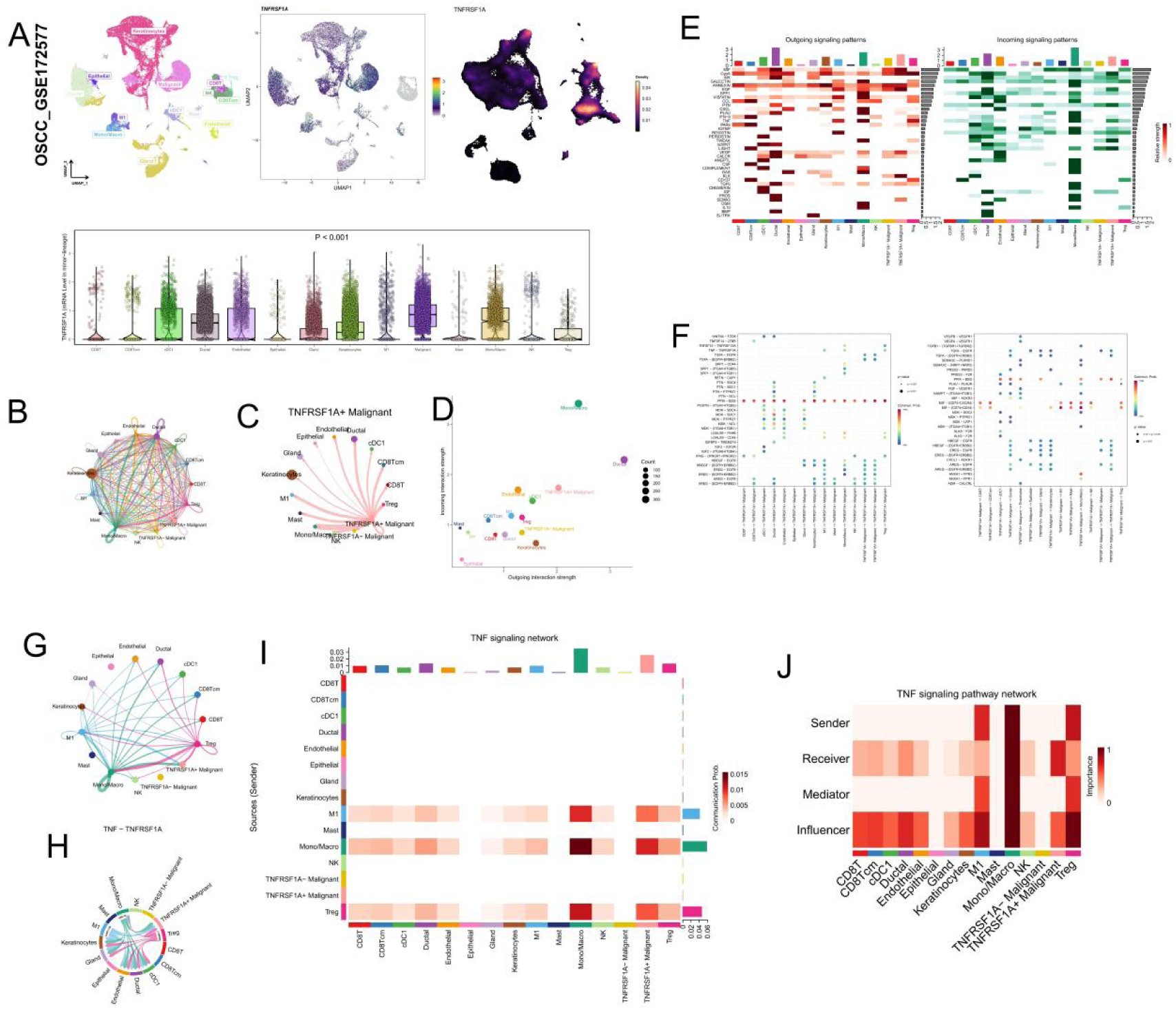
Single-cell communication networks of TNFRSF1A signaling in OSCC. (A) UMAP visualization of OSCC (GSE117257) single-cell data showing TNFRSF1A expression across cell clusters. (B–C) Intercellular communication networks of TNFRSF1A⁺ malignant cells with other cell populations. (D) Scatter plots showing the overall outgoing signaling strength of each cell type. (E–F) Heatmaps of outgoing and incoming signaling patterns across cell populations. (G–H) Network diagrams of TNF– TNFRSF1A signaling interactions. (I–J) TNF signaling network analysis showing sender, receiver, mediator, and influencer roles for each cell type, highlighting TNFRSF1A⁺ malignant cells as major signal transducers.

## Discussion

*F. nucleatum*, an anaerobic gram-negative bacterium, has traditionally been studied as a periodontal pathogen[21–23]. However, recent research has expanded its relevance to tumor biology[24–28]. *F. nucleatum* has been detected in biopsy specimens from patients with colorectal and other malignancies, where its presence correlates with increased tumor invasiveness and poor prognosis[29, 30].

Both *F. nucleatum* and other gram-negative bacteria release outer membrane vesicles (OMVs) *in vitro* and *in vivo*. These nanoparticles, typically ranging from 50 to 250 nm in diameter, play a critical role in bacterial pathogenicity. OMVs often contain lipopolysaccharides (LPS), DNA, adhesins, and enzymes[28, 31], functioning as carriers for these virulent factors[32, 33]. In cancer research, OMVs are considered pivotal in the tumor microenvironment, where they enhance tumor cell proliferation, invasion, and metastasis, and modulate host immune responses[12, 13]. It has been demonstrated that *F. nucleatum* produces OMVs[34], yet the link between *F. nucleatum* OMVs and carcinogenesis remains to be fully elucidated.

*F. nucleatum* promotes tumor development through the NF-κB signaling cascade, a well-established pathway in the progression of various cancers, including oral cancer[35]. NF-κB is activated *via* two main pathways: the classical and alternative routes[36]. The classical pathway begins with the activation of tumor necrosis factor (TNF) receptor-associated factor (TRAF) adapter proteins, leading to the stimulation of the IκB kinase (IKK) complex, which comprises IKK1, IKK2, and the structural/regulatory component NF-κB essential modulator (NEMO) (Figure 7). The active IKK complex phosphorylates IκB proteins, targeting them for proteasomal degradation. This process liberates NF-κB dimers, enabling their translocation to the nucleus. A series of experiments were conducted to investigate the effects of *F. nucleatum* OMVs on TNFR1 and NF-κB signaling pathways in OSCC cells. A multidimensional analysis approach was employed, including transmission electron microscopy, nanoparticle size analysis, transcriptome sequencing, and both *in vitro* and *in vivo* experiments. The results demonstrated that *F. nucleatum* and its OMVs play a significant role in promoting the proliferation and invasion of OSCC cell lines CAL27 and SCC25, primarily through the TNFR1 and NF-κB signaling pathways. In contrast, *Porphyromonas gingivalis* (*Pg*) did not significantly affect TNFR1 protein expression under the same conditions, highlighting the unique role of *F. nucleatum* in carcinogenesis.

Our findings demonstrate that FnOMVs induced greater proliferation in OSCC cells and led to the development of larger tumors *in vivo* compared to Fn, indicating a potentially stronger promotive effect on tumor growth. The vesicle membrane structure of OMVs may shield pathogenic agents from degradation, enabling their concentrated delivery to host cells *via* endocytosis, which facilitates the transmission of harmful components and the activation of proinflammatory or necrotic pathways[37, 38]. This suggests that FnOMVs may exhibit greater carcinogenic potential than the bacterium itself. Additionally, FnOMVs have been shown to stimulate the epithelial-mesenchymal transition (EMT) phenotype in OSCC cells, enhancing cell invasion and migration in a time- and concentration-dependent manner[28]. These findings not only complement existing research but also offer a new perspective on how oral microbiota influence cancer cells through indirect mechanisms, further confirming the critical role of oral microbiota in OSCC development.

Despite the advances made in understanding the roles of *F. nucleatum* and OMVs in OSCC progression, this study has certain limitations. The specific components within OMVs responsible for activating TNFR1 and promoting proliferation and invasion have not yet been identified. Future research should focus on isolating these active components and elucidating their interaction mechanisms with host cell receptors. This study not only deepens our understanding of the interaction mechanisms between oral microbiota and OSCC but also lays the groundwork for novel therapeutic strategies targeting oral microbiota. Targeting OMVs as therapeutic agents offers a promising new approach to inhibiting the tumor-promoting effects of *F. nucleatum* on OSCC. Furthermore, this research underscores the importance of considering microbial factors in cancer therapy, paving the way for advancements in precision medicine.

**Figure 7.**
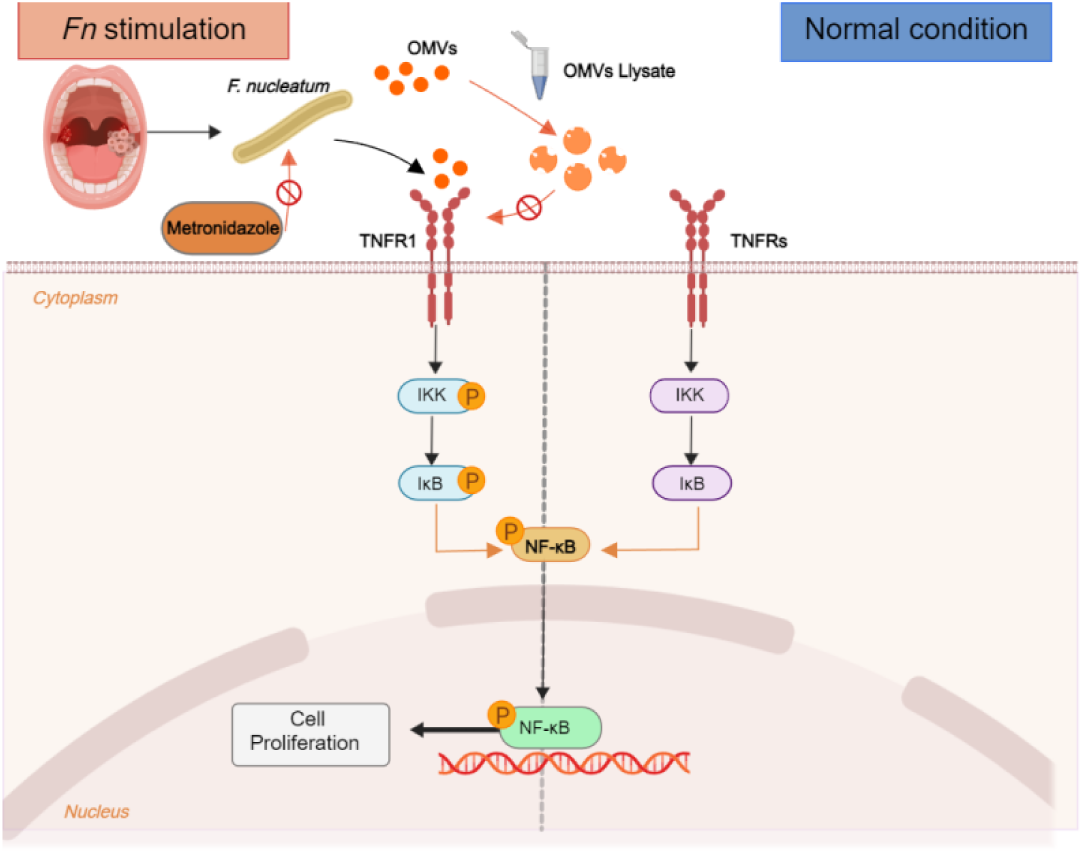
Fn OMVs Activate the NF-κB Signaling Pathway *via* TNFR1 to Promote OSCC Cell Proliferation.

## Conclusion

This study highlights the critical role of *Fusobacterium nucleatum* and its OMVs in driving OSCC proliferation *via* the TNFR1/NF-κB signaling pathway. The findings highlight a higher prevalence of *F. nucleatum* in OSCC tissues and demonstrate that both the bacterium and its OMVs significantly enhance OSCC cell growth and proliferation. *In vivo*, OMVs from *F. nucleatum* led to the development of larger tumors compared to the bacterium alone, emphasizing their increased tumorigenic potential. This research deepens the understanding of microbial influences on cancer progression, pointing to potential therapeutic targets within microbial regulatory mechanisms. Future studies should aim to identify the specific OMV components responsible for these effects and to validate these findings clinically. Although there are limitations, such as the need to identify precise molecular components, this study provides a crucial foundation that could shape clinical practices and policy development in cancer treatment by integrating the microbiome’s role.

## CRediT author statement

Zhenrui Li: Conceptualization, Data curation, Formal analysis, Methodology, Visualization, Writing–original draft; Ji’an Liu: Conceptualization, Data curation, Methodology, Writing–original draft; Rao Fu: Formal analysis, Visualization, Writing–original draft; Xutao Wen: Formal analysis, Software, Visualization; Xufeng Huang: Formal analysis, Investigation, Resources; Divya Gopinath: Project administration, Supervision, Writing–review & editing; Ling Zhang: Funding acquisition, Investigation, Project administration,Supervision, Writing–review & editing.

## Reference

1. Sung H, Ferlay J, Siegel RL, Laversanne M, Soerjomataram I, Jemal A, et al. Global Cancer Statistics 2020: GLOBOCAN Estimates of Incidence and Mortality Worldwide for 36 Cancers in 185 Countries. CA Cancer J Clin. 2021; 71: 209–49.

2. Ferlay J, Colombet M, Soerjomataram I, Parkin DM, Piñeros M, Znaor A, et al. Cancer statistics for the year 2020: An overview. Int J Cancer. 2021.

3. Li Z, Liu Y, Zhang L. Role of the microbiome in oral cancer occurrence, progression and therapy. Microb Pathog. 2022; 169: 105638.

4. Gopinath D, Menon RK, Banerjee M, Su Yuxiong R, Botelho MG, Johnson NW. Culture-independent studies on bacterial dysbiosis in oral and oropharyngeal squamous cell carcinoma: A systematic review. Crit Rev Oncol Hematol. 2019; 139: 31–40.

5. Brennan CA, Garrett WS. Fusobacterium nucleatum - symbiont, opportunist and oncobacterium. Nat Rev Microbiol. 2019; 17: 156–66.

6. Cai L, Zhu H, Mou Q, Wong PY, Lan L, Ng CWK, et al. Integrative analysis reveals associations between oral microbiota dysbiosis and host genetic and epigenetic aberrations in oral cavity squamous cell carcinoma. NPJ Biofilms Microbiomes. 2024; 10: 39.

7. Saikia PJ, Pathak L, Mitra S, Das B. The emerging role of oral microbiota in oral cancer initiation, progression and stemness. Front Immunol. 2023; 14: 1198269.

8. Li Z, Liu Y, Huang X, Wang Q, Fu R, Wen X, et al. F. Nucleatum enhances oral squamous cell carcinoma proliferation via E-cadherin/β-Catenin pathway. BMC Oral Health. 2024; 24: 518.

9. Li L, Chandra V, McAllister F. Tumor-resident microbes: the new kids on the microenvironment block. Trends Cancer. 2024; 10: 347–55.

10. Lu YQ, Qiao H, Tan XR, Liu N. Broadening oncological boundaries: the intratumoral microbiota. Trends Microbiol. 2024.

11. Despins CA, Brown SD, Robinson AV, Mungall AJ, Allen-Vercoe E, Holt RA. Modulation of the Host Cell Transcriptome and Epigenome by Fusobacterium nucleatum. mBio. 2021; 12: e0206221.

12. Dell’Annunziata F, Folliero V, Giugliano R, De Filippis A, Santarcangelo C, Izzo V, et al. Gene Transfer Potential of Outer Membrane Vesicles of Gram-Negative Bacteria. Int J Mol Sci. 2021; 22.

13. Cui C, Guo T, Zhang S, Yang M, Cheng J, Wang J, et al. Bacteria-derived outer membrane vesicles engineered with over-expressed pre-miRNA as delivery nanocarriers for cancer therapy. Nanomedicine. 2022; 45: 102585.

14. Guo Q, Jin Y, Chen X, Ye X, Shen X, Lin M, et al. NF-κB in biology and targeted therapy: new insights and translational implications. Signal Transduct Target Ther. 2024; 9: 53.

15. Ebrahimi N, Abdulwahid ARR, Mansouri A, Karimi N, Bostani RJ, Beiranvand S, et al. Targeting the NF-κB pathway as a potential regulator of immune checkpoints in cancer immunotherapy. Cell Mol Life Sci. 2024; 81: 106.

16. Oeckinghaus A, Ghosh S. The NF-kappaB family of transcription factors and its regulation. Cold Spring Harb Perspect Biol. 2009; 1: a000034.

17. Jackson-Bernitsas DG, Ichikawa H, Takada Y, Myers JN, Lin XL, Darnay BG, et al. Evidence that TNF-TNFR1-TRADD-TRAF2-RIP-TAK1-IKK pathway mediates constitutive NF-kappaB activation and proliferation in human head and neck squamous cell carcinoma. Oncogene. 2007; 26: 1385–97.

18. Zhang Z, Sun D, Tang H, Ren J, Yin S, Yang K. PER2 binding to HSP90 enhances immune response against oral squamous cell carcinoma by inhibiting IKK/NF-κB pathway and PD-L1 expression. J Immunother Cancer. 2023; 11.

19. Chiu HW, Lee HL, Lee HH, Lu HW, Lin KY, Lin YF, et al. AIM2 promotes irradiation resistance, migration ability and PD-L1 expression through STAT1/NF-κB activation in oral squamous cell carcinoma. J Transl Med. 2024; 22: 13.

20. Dan H, Liu S, Liu J, Liu D, Yin F, Wei Z, et al. RACK1 promotes cancer progression by increasing the M2/M1 macrophage ratio via the NF-κB pathway in oral squamous cell carcinoma. Mol Oncol. 2020; 14: 795–807.

21. Han YW. Fusobacterium nucleatum: a commensal-turned pathogen. Curr Opin Microbiol. 2015; 23: 141–7.

22. Chen Y, Huang Z, Tang Z, Huang Y, Huang M, Liu H, et al. More Than Just a Periodontal Pathogen -the Research Progress on Fusobacterium nucleatum. Front Cell Infect Microbiol. 2022; 12: 815318.

23. Wang Y, Wang L, Sun T, Shen S, Li Z, Ma X, et al. Study of the inflammatory activating process in the early stage of Fusobacterium nucleatum infected PDLSCs. Int J Oral Sci. 2023; 15: 8.

24. Rubinstein MR, Wang X, Liu W, Hao Y, Cai G, Han YW. Fusobacterium nucleatum promotes colorectal carcinogenesis by modulating E-cadherin/β-catenin signaling via its FadA adhesin. Cell Host Microbe. 2013; 14: 195–206.

25. Zhu H, Li M, Bi D, Yang H, Gao Y, Song F, et al. Fusobacterium nucleatum promotes tumor progression in KRAS p.G12D-mutant colorectal cancer by binding to DHX15. Nat Commun. 2024; 15: 1688.

26. Ponath F, Tawk C, Zhu Y, Barquist L, Faber F, Vogel J. RNA landscape of the emerging cancer-associated microbe Fusobacterium nucleatum. Nat Microbiol. 2021; 6: 1007–20.

27. Parhi L, Alon-Maimon T, Sol A, Nejman D, Shhadeh A, Fainsod-Levi T, et al. Breast cancer colonization by Fusobacterium nucleatum accelerates tumor growth and metastatic progression. Nat Commun. 2020; 11: 3259.

28. Chen G, Gao C, Jiang S, Cai Q, Li R, Sun Q, et al. Fusobacterium nucleatum outer membrane vesicles activate autophagy to promote oral cancer metastasis. J Adv Res. 2024; 56: 167–79.

29. Gopinath D, Menon RK, Wie CC, Banerjee M, Panda S, Mandal D, et al. Differences in the bacteriome of swab, saliva, and tissue biopsies in oral cancer. Sci Rep. 2021; 11: 1181.

30. Neuzillet C, Marchais M, Vacher S, Hilmi M, Schnitzler A, Meseure D, et al. Prognostic value of intratumoral Fusobacterium nucleatum and association with immune-related gene expression in oral squamous cell carcinoma patients. Sci Rep. 2021; 11: 7870.

31. Pignatelli P, Nuccio F, Piattelli A, Curia MC. The Role of Fusobacterium nucleatum in Oral and Colorectal Carcinogenesis. Microorganisms. 2023; 11.

32. Toyofuku M, Nomura N, Eberl L. Types and origins of bacterial membrane vesicles. Nat Rev Microbiol. 2019; 17: 13–24.

33. Sartorio MG, Pardue EJ, Feldman MF, Haurat MF. Bacterial Outer Membrane Vesicles: From Discovery to Applications. Annu Rev Microbiol. 2021; 75: 609–30.

34. Liu J, Hsieh CL, Gelincik O, Devolder B, Sei S, Zhang S, et al. Proteomic characterization of outer membrane vesicles from gut mucosa-derived fusobacterium nucleatum. J Proteomics. 2019; 195: 125–37.

35. Nakayama H, Ikebe T, Beppu M, Shirasuna K. High expression levels of nuclear factor kappaB, IkappaB kinase alpha and Akt kinase in squamous cell carcinoma of the oral cavity. Cancer. 2001; 92: 3037–44.

36. Hayden MS, Ghosh S. Shared principles in NF-kappaB signaling. Cell. 2008; 132: 344–62.

37. Engevik MA, Danhof HA, Ruan W, Engevik AC, Chang-Graham AL, Engevik KA, et al. Fusobacterium nucleatum Secretes Outer Membrane Vesicles and Promotes Intestinal Inflammation. mBio. 2021; 12.

38. Briaud P, Carroll RK. Extracellular Vesicle Biogenesis and Functions in Gram-Positive Bacteria. Infect Immun. 2020; 88.

